# Benchmarking AI-generated structural ensembles of membrane proteins against physics-based modelling

**DOI:** 10.64898/2026.08.08.743655

**Authors:** Bryony RJ Clifton, Adam G Grieve, Robin A Corey

## Abstract

Proteins dynamically switch between a continuum of interconverting conformational states, and understanding these structural dynamics is important for understanding protein function and for developing therapeutics. Molecular dynamics (MD) simulations can provide insight into protein conformational ensembles, but sampling rare conformational states can require substantial computational resources. The recent development of AI-based approaches for generating protein conformational ensembles, such as the Biomolecular Emulator (BioEmu), offers a potential alternative, although it remains unclear whether these approaches can accurately capture the conformational landscapes, especially for special cases such as membrane proteins. Here, we assess the ability of BioEmu to model the conformational dynamics of a model membrane protein, the bacterial rhomboid intramembrane proteases GlpG. We find that BioEmu generates a range of conformations corresponding to both open and closed states of the rhomboid lateral gate, including states associated with different stages of the catalytic cycle. These conformations broadly correspond to states sampled during microsecond-timescale MD simulations, although BioEmu does not reproduce the full conformational landscape observed using MD. BioEmu also samples substantial conformational heterogeneity within the soluble domains of rhomboids, which are highly flexible and poorly represented in experimental structures. Overall, our findings demonstrate that BioEmu can generate plausible conformational ensembles for relatively large, six-and seven-pass membrane proteins, sampling rare states at a fraction of the computational cost of conventional MD simulations. These results suggest that AI-based ensemble generation could provide an accessible approach for exploring membrane protein dynamics and complement conventional molecular modelling approaches.

## Introduction

Proteins do not exist in a single static conformation but instead dynamically switch within a continuum of interconverting microstates. Understanding these structural dynamics is crucial for understanding protein function and for designing therapeutics that modulate protein activity. However, there are many associated challenges, not least the difficulty of experimentally determining complete protein conformational ensembles. Approaches such as time-resolved cryo-EM, which trap conformational states at different time points following addition of ligand, or at different ligand concentrations, can facilitate the identification of transient states (*1, 2*). However, an additional level of complexity is introduced when working with membrane proteins. Membrane proteins are especially challenging, partly due to their hydrophobicity, which makes them difficult to purify. Attempts to ameliorate this, such as through reconstitution into non-native detergents or artificial lipid particles, can dramatically alter structure and function.

Molecular modelling approaches, such as molecular dynamics (MD), offer a powerful tool for overcoming some of these challenges. By simulating proteins within physiologically relevant membrane environments, MD can sample conformational ensembles and estimate the relative free energies of conformational transitions. However, some conformational changes are associated with high energy barriers, which mean that they happen very rarely in conventional equilibrium simulations, requiring advanced computational resources and long runtimes. Enhanced sampling methods can be applied to identify rarer states and, whilst these may overcome some issues, they can be challenging to set up and converge, are susceptible to sampling bias or other technical artefacts, and can still fail to capture all possible states. Perhaps most importantly, these modelling techniques rely on high-quality input structures. As many membrane proteins still lack experimentally determined high-resolution structures, the application of molecular dynamics has historically been limited by the availability of suitable starting models(*3*).

The development of AI-based structure prediction tools, such as AlphaFold 2 and 3, has led to an incredible advance in our understanding of protein function(*4, 5*). Many of these tools show an impressive level of precision and accuracy in predicting previously unsolved structures, opening up potential for the advanced study of otherwise neglected proteins. Whilst AlphaFold does not directly include the membrane environment during the modelling process, it has demonstrated moderately high accuracy modelling of membrane proteins, likely due to the presence of membrane proteins in its training dataset(*6, 7*).

Whilst AlphaFold has shown a modest ability to predict the dynamics of membrane proteins, it typically only predicts a limited range of microstates. Specific conformations are often omitted, such as closed ion channels or active receptors, unless adaptations or a variety of AI-based approaches are applied(*8–10*). This limits the functional insights we can make into proteins from structure prediction alone. Therefore, MD is often required to gain insights into a wider conformational landscape.

The Biomolecular Emulator, or BioEmu, offers a novel approach for generating conformational ensembles of proteins (*11*). Built upon concepts from the AlphaFold2 Evoformer architecture and further trained using MD trajectories together with structurally heterogeneous entries from the AlphaFold Protein Structure Database (AFDB), BioEmu has been reported to generate conformational ensembles that approximate the equilibrium distributions of small- to medium-sized soluble proteins. In benchmark studies, BioEmu has reproduced conformational states that are difficult to sample using conventional MD simulations, whilst requiring only a fraction of the computational cost(*11, 12*).

However, it is currently unclear whether BioEmu can accurately model the conformational landscapes of membrane proteins, as these were largely absent from the model or training data(*11*). Here, we present a case study in which BioEmu is applied to the rhomboid family of intramembrane proteases, to assess its ability to predict structural dynamics of membrane proteins.

Rhomboids are essential and ubiquitous enzymes, known to regulate signalling and quality control(*13–18*). The bacterial rhomboid, GlpG, has been demonstrated to exist in open and closed states, with opening of the lateral gate, formed between transmembrane helices (TM) 2 and 5, required for its proteolytic activity(*19–23*). This is also considered to be the case for human rhomboids, as demonstrated by MD simulations of AlphaFold2 models. Without MD, these AlphaFold2, as well as Boltz-2, and Chai-1 models displayed very limited conformational heterogeneity (*24*).

Additionally, all human rhomboids and *E. coli* GlpG feature a soluble cytoplasmic (or inner mitochondrial membrane) domain, which is thought to function independent of the core proteolytic activity of the enzyme. Although the structure of the GlpG soluble domain has been solved in isolation, it is excluded from X-ray crystal structures that include the core GlpG transmembrane region, due to its high flexibility(*22, 25, 26*). How the soluble domains of rhomboids regulate their activity largely remains unclear, with few studied examples of their role(*27, 28*).

In this study, we evaluate the ability of BioEmu to generate conformational ensembles for rhomboid proteases spanning both open and closed conformations. We observe that BioEmu recapitulates a variety of open and closed states, corresponding to conformations associated with different stages of the catalytic cycle, as sampled in µs-long MD simulations. Although it does not recreate all states seen via MD modelling, we show that BioEmu allows sampling of rare conformational states of relatively large, six- and seven-pass membrane proteins using only a fraction of the computational time and resources required for conventional MD simulations. These findings suggest that BioEmu could make studies of membrane protein dynamics more accessible whilst substantially reducing computational costs.

## Results

### BioEmu lacks lipid bilayer context for prediction of membrane protein structures

Of the rhomboid proteases, experimental structures have only been solved for bacterial rhomboids, with the GlpG protease being the best characterised, with both open and closed states identified(*22, 26*). However, these structures only comprise the core, 6-pass transmembrane region, with the flexible cytoplasmic domain not modelled. Therefore, we sought both to test the ability of BioEmu to capture biologically relevant conformational dynamics of the GlpG rhomboid fold, as well as to investigate the conformational heterogeneity predicted for the soluble domain.

Of 1000 GlpG samples predicted by BioEmu, a sampling scale in line with previous studies(*12, 29*), 969 passed filtering for unphysical structures. To assess the conformational distribution of these samples, FoldSeek was used to cluster the models, using a template modelling threshold of 0.7 to account for the identical sequence used across the query (*30*). Four clusters were obtained, as shown in Figure 1. Cluster 1 was predominant, containing 936 models, whereas clusters 2-4 contained 29, 3, and 1 models respectively.

**Figure 1.**
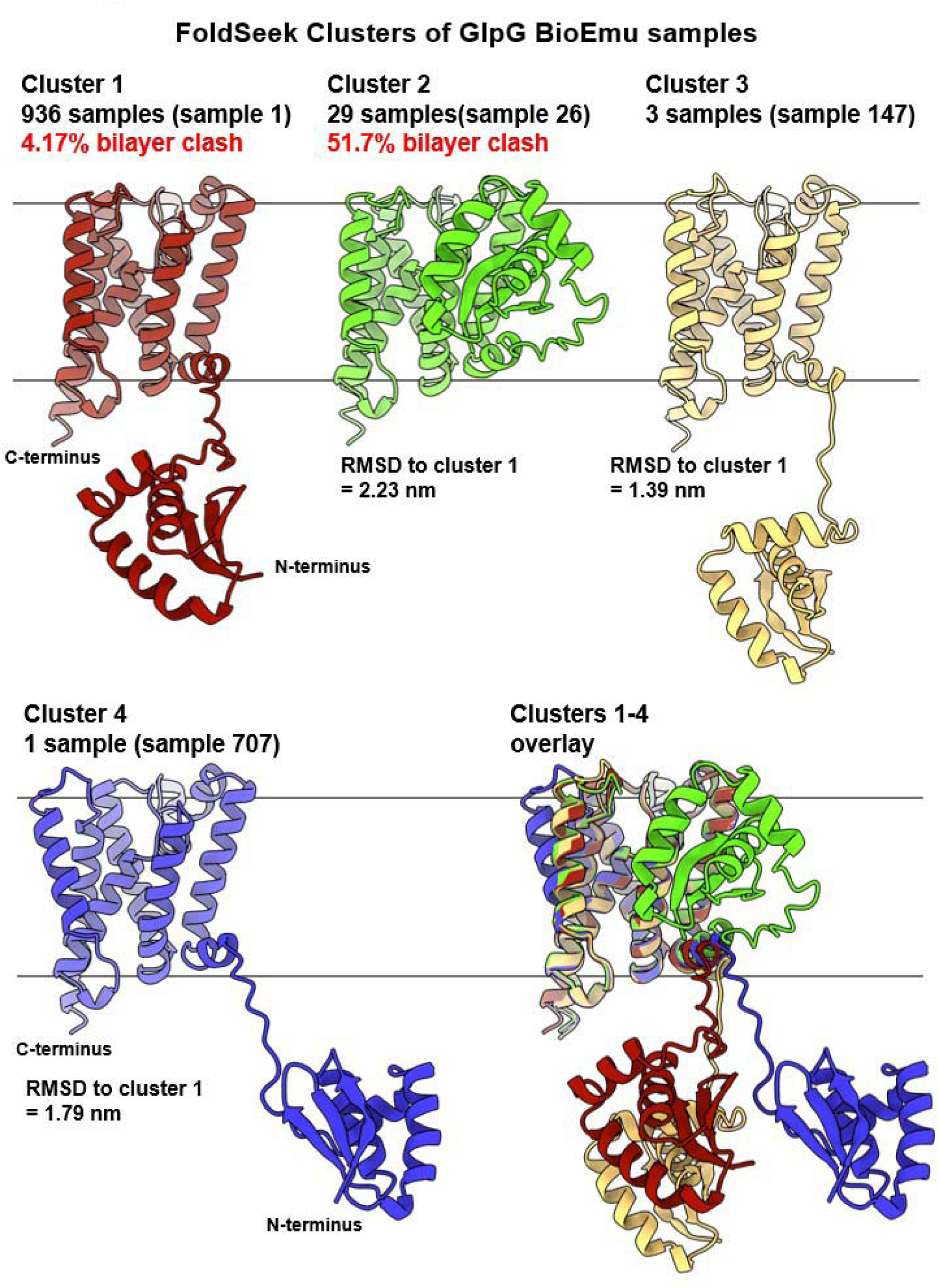
FoldSeek clustering was applied to BioEmu predictions of GlpG, using a template modelling threshold of 0.7. The representative model for each cluster is depicted by a cartoon representation. The number of models in each cluster is indicated and the approximate lipid bilayer position is depicted by grey lines.

The structure of the GlpG soluble domain was highly consistent across all models, with only one notable outlier when compared to a previously-published structure (Figure S1)(*25*). Aligning the representative sample from each cluster shows that, for the first 3 clusters, positioning of the soluble domain relative to the core transmembrane fold was the primary source of heterogeneity. For the representative sample from cluster 2, the soluble domain lay within the predicted plane of the lipid bilayer and TM core of GlpG, which is extremely unlikely in a physiological system.

To determine how many of the 969 models feature soluble domains in physically implausible positions, all BioEmu samples were aligned against an AlphaFold prediction embedded within a model lipid bilayer using PPM 2.0(*31*). The soluble domain in 54 total models (5.6%) was found to penetrate deeper than the 0.5 nm polar region that would be typical of a minor insertion and these are therefore considered physically implausible. 39 of these correspond to cluster 1 models (4.17% of samples) and 15 to cluster 2 (51.7% of samples)(Figure 1). A further 57 (5.9%) models featured small sections of the soluble domain within the polar region of the lipid bilayer and were therefore considered within reason for a physiological system (see Figure S2 for examples of plausible and implausible models).

### BioEmu successfully captures structural variance at the lateral gate of the bacterial rhomboid GlpG

RMSF analysis was performed to identify specific regions which were the most variable between the BioEmu models (Figure S3). As expected, the soluble domain exhibited the greatest variability across the 969 samples. As the variability in this region is so high compared to that of the core fold, it likely dominates in the clustering, preventing relatively smaller differences within the core fold from emerging. This could explain why, in the full-length clusters, the only difference within the core fold is seen for cluster 4, which consisted of only 1 sample (sample 707). In sample 707, a dramatic difference in the positioning of both L5 and TM5, with a dramatic outward movement of TM5 is exhibited (Figure 1). Relatively minor differences between samples from each cluster were also seen in loop 5 (L5), connecting TM5 and TM6.

To more closely examine the specific regions within the core fold that differ across the predictions, analyses were repeated with the soluble N-terminus discounted (GlpG ΔN). RMSF heatmaps of GlpG ΔN demonstrated that the hotspots for deviation across models are the few residues at the very C-terminal end of TM6; TM5; the L5 and L6 loops connecting TM5 to TM4/6; and the L1 loop (Figure S3). TM5 movement via flexibility of the L5 loop has been previously demonstrated to be required for proteolysis by GlpG, and its outward movement defines the ‘gate open’ and ‘gate closed’ states(*21, 26*). The flexibility in this region captured across different BioEmu samples suggests that the structure prediction tool can capture different conformational states which are important for the molecular function of GlpG.

To further investigate the different conformations of the GlpG core fold which were predicted by BioEmu, FoldSeek clustering was repeated for GlpG ΔN. To obtain more than 1 cluster containing multiple samples, a template modelling threshold of 0.98 was required, reflecting the high structural similarity across these models. Using these parameters, 10 clusters were obtained (Figure S4). Clusters 1 and 2 comprised 957 and 4 samples respectively, with all others comprising a single sample.

To investigate whether the different GlpG ΔN clusters represent models with diverse lateral gate conformations, lateral gate quantification was performed for all 969 samples as previously described (Figure S5A)(*24*). This analysis revealed a diverse landscape of lateral gate conformations, with 12.8% of models occupying an open state, previously defined as ∼1.4 nm lateral gate distance between TM2 and TM5(*24*). The median lateral gate distance from BioEmu models was 1.35 nm, with the majority of models distributed around this value (interquartile range or IQR = 1.32-1.37 nm). However, minimum and maximum values of 1.16 nm and 2.02 nm were observed, demonstrating an ability of BioEmu to predict extreme conformational states.

Splitting the lateral gate distances by cluster demonstrates that cluster 1 primarily represents models around this median value of lateral gate distance, with fully open or closed states mostly seen in the other clusters (Figure S5B). For instance, cluster 2 models are all in an open state, as can be seen by the L5 loop lifting away from the extracellular face of GlpG and TM5 moving outwards (Figure 2A-C). Conversely, the cluster 8 model represents a closed state, with TM5 closer to TM2 and the L5 loop fully occluding access to the active site (Figure 3A-C). Overall, these data demonstrate that BioEmu can successfully model both open and closed states of the GlpG protease.

**Figure 2.**
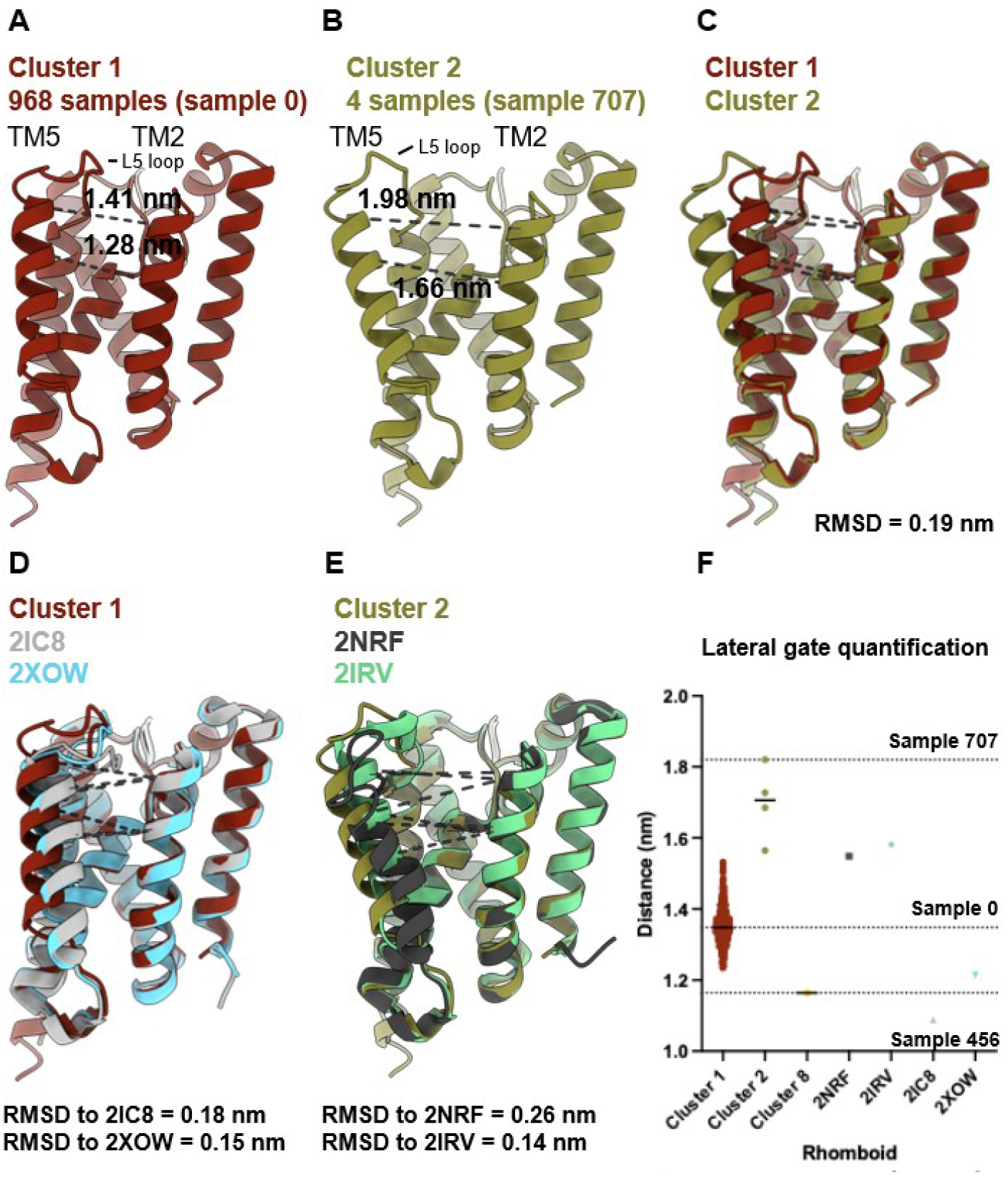
Atoms from residues Met1 to Ala93 were removed to form GlpG ΔN. FoldSeek clustering was then applied, using a template modelling threshold of 0.98. The representative models of **(A)** cluster 1 and **(B)** cluster 2 are depicted as cartoons, with lateral gate quantification displayed by dashed lines and values. **(C-E)** Structural alignment of clusters 1 and 2 with each other, and GlpG crystal structures. RMSD values to the BioEmu model are displayed below. **(F)** Lateral gate quantification of BioEmu models, stratified by clusters which represent the largest differences in lateral gate distance (clusters 1, 2, and 8), and crystal structures. Solid line = median lateral gate distance; dashed lines = lateral gate distance of sample 0, 456, or 707 as labelled.

**Figure 3.**
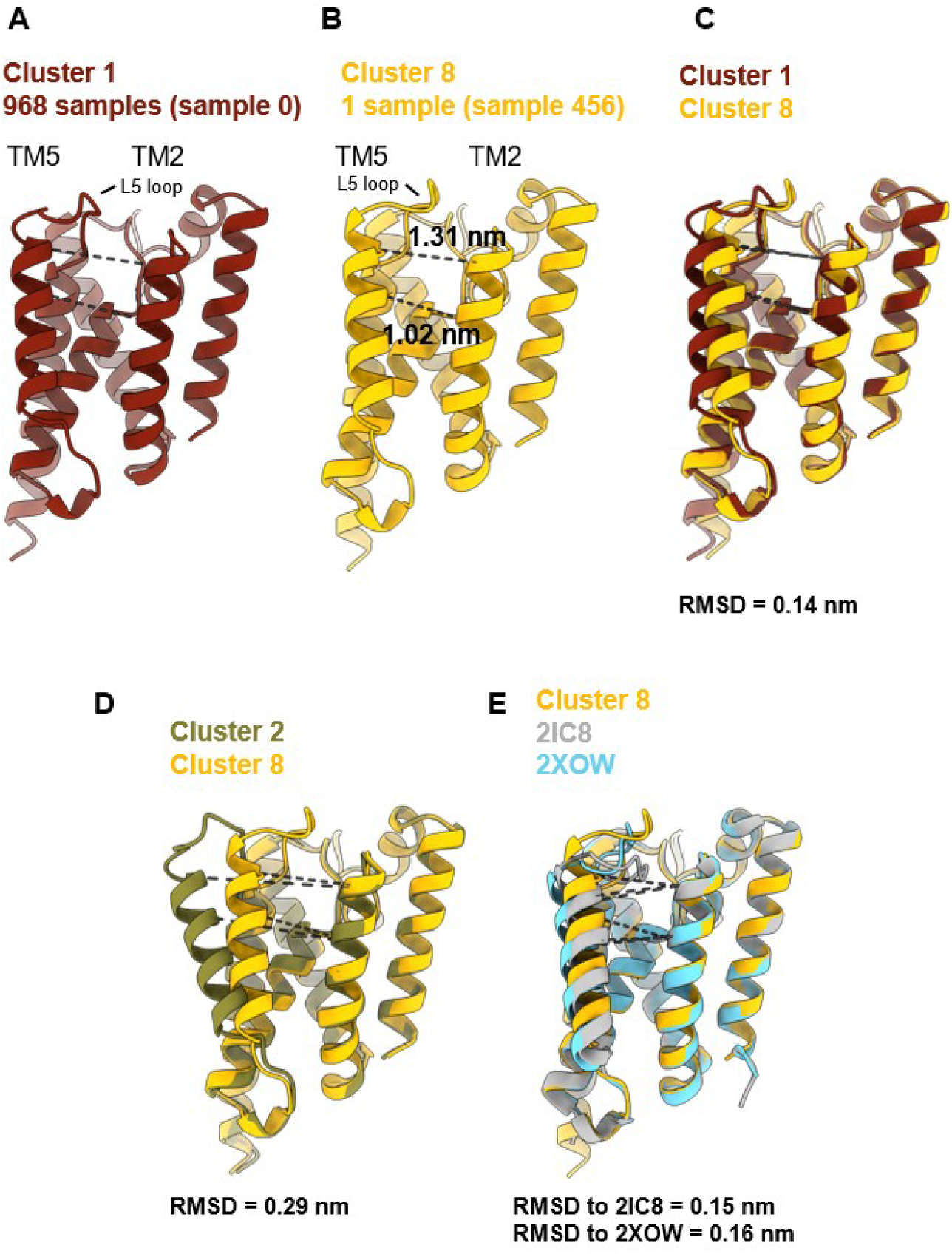
Cartoon depictions of representative models for **(A)** cluster 1 and **(B)** cluster 8, obtained from FoldSeek clustering of GlpG ΔN. Dashed lines represent positioning of atoms used for lateral gate quantification. **(C-E)** Structural alignments of cluster 8, with RMSD displayed below.

### BioEmu generates states closely resembling experimental models in addition to unique states

To examine how well these BioEmu predicted structures correspond to previously generated experimental models, structural alignments with crystal structures were performed, revealing high conservation of the core fold. Overlays from alignments with closed structures (2IC8 and 2XOW) revealed that the cluster 1 representative model, sample 0, occupies a partially open state (Figure 2D). The lateral gate for this model is wider than either of the X-ray crystal structures (sample 0 = 1.34 nm; 2IC8 = 1.09 nm; 2XOW = 1.21 nm). Furthermore, the L5 loop is comparatively lifted in the cluster 1 model, leaving the active site more accessible than in either of the crystal structures.

Cluster 2 comprises 4 models, with lateral gate distances ranging from 1.56 nm to 1.82 nm (Figure 2E). Alignments with open structures (2NRF, chain A and 2IRV, chain B) reveal that the cluster 2 representative model, sample 707, is highly similar to 2IRV. The L5 loop is lifted away from the core fold and the lateral gate is fully open. Remarkably, the lateral gate is even wider than for either of the open crystal structures (sample 707 = 1.82 nm; 2NRF = 1.55 nm; 2IRV = 1.58 nm), which suggests that BioEmu can predict a diversity of states, including those which may not be readily captured by structural biology approaches.

Interestingly, the monotypic cluster 8 model, sample 456, is the only 1 of 969 models which possess a lateral gate as narrow as the closed crystal structures 2IC8 and 2XOW (Figure 2F). This model features a lateral gate distance of 1.16 nm (Figure 3B). Structural overlays demonstrate that this model is highly similar to that of both the inhibitor bound and apo closed structures of GlpG (Figure 3E).

To further investigate how much the different lateral gate conformations correspond to variation between BioEmu GlpG ΔN models, the RMSD for each sample against different reference structures or models were plotted (Figure S5C). The resulting scatter plots demonstrated that the samples which deviate most from the other models are within clusters 2-10, supporting the previous conclusion that the clusters primarily group by different lateral gate states.

Overall, our data demonstrate that BioEmu can accurately predict GlpG models that have a variety of biologically relevant lateral gate states, including those seen in experimental structures in addition to unique conformational states not previously reported.

### BioEmu can predict conformational ensembles of membrane proteins but under-samples relative to molecular dynamics simulations

We next ran principal component analysis on the BioEmu GlpG ΔN models to assess the range of structural sampling of the BioEmu data in relation to ensembles generated using molecular dynamics (MD) simulations.

First, PCA was performed on the BioEmu sampling alone (Figure 4A-B, S6). Projecting eigenvector 1, which chiefly describes the outward movement of TM5 (Figure 4A), reveals a similar distribution to that seen by the earlier lateral gate quantification, with the tails represented almost entirely by FoldSeek clusters 2-10 (Figures 4B, S5B and S6A-B). Additionally, many BioEmu predictions overlap with the closed crystal structures, whereas there is less overlap with the open structures (Figure 4B). However, it is surprising that there are not distinct clusters within the continuum of states, as might be expected for a protein with multiple distinct functional states.

**Figure 4.**
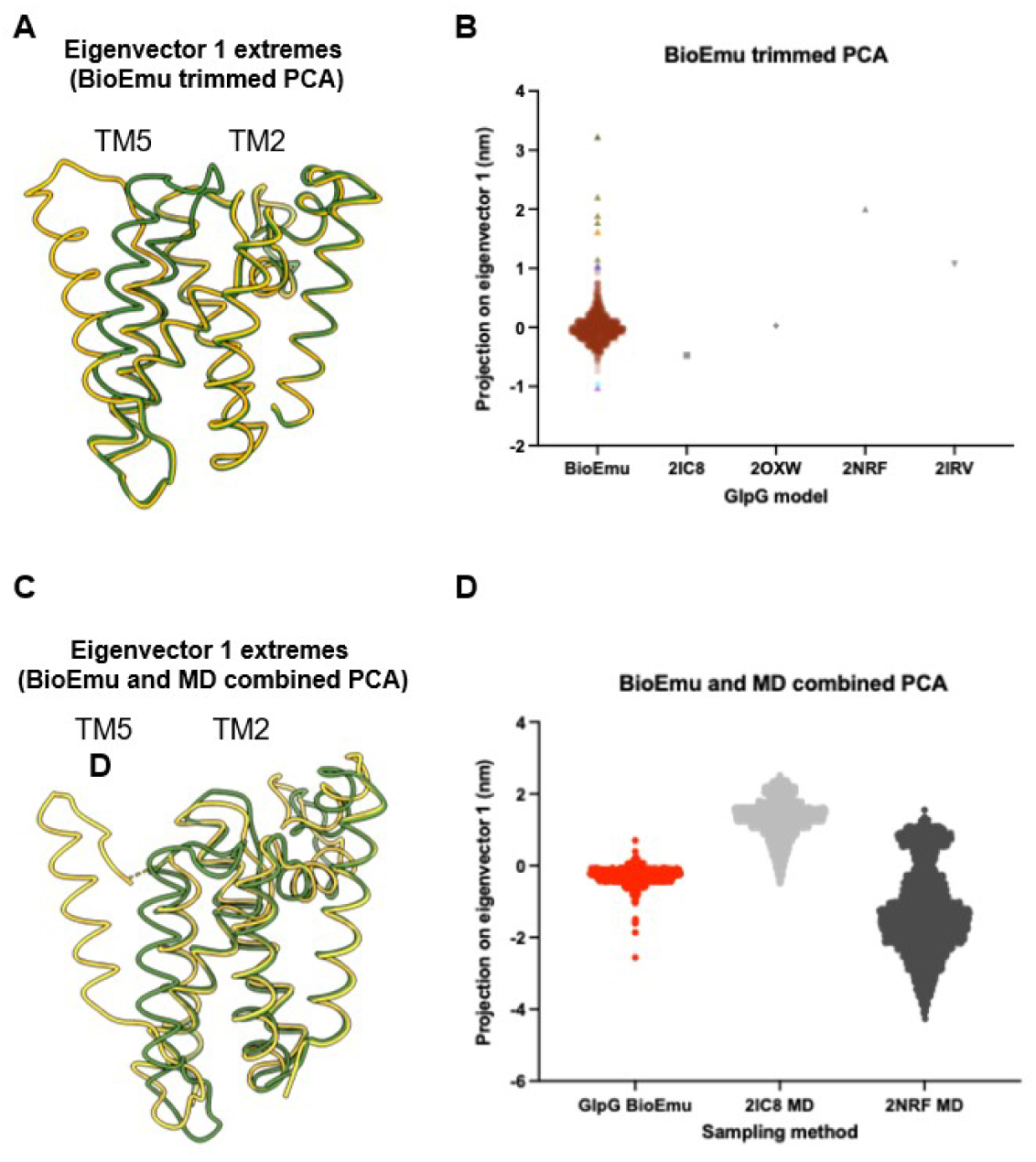
Principal component analysis (PCA) was performed using GlpG residues Ala93 to Leu270 (‘GlpG trimmed’). **(A)** Overlay of extreme models for eigenvector 1 (27.6% of total variance) of GlpG BioEmu PCA. **(B)** All GlpG samples were projected onto eigenvector 1 and visualised by cluster. All cluster 1 samples are shown as translucent red circles. All other clusters are opaque triangles, with colours matching those in Figure S2. GlpG crystal structures were also projected onto eigenvector 1, as displayed by grey shapes. **(C)** Overlay of extreme models for eigenvector 1 (28.9% total variance) of a combined PCA for GlpG BioEmu samples and MD simulations for 2IC8 and 2NRF, as previously reported(*24*). **(D)** All GlpG BioEmu and MD data were projected onto eigenvector 1 and stratified by sampling method.

Our analyses so far have demonstrated that BioEmu can accurately report an ensemble of biologically relevant conformational states for GlpG. However, it has been reported that BioEmu can match, or even surpass, the ability of MD to sample structural ensembles for soluble proteins. To assess whether this is also true for membrane proteins, the GlpG BioEmu predictions were benchmarked against a previously published MD dataset for GlpG, which also lacked a soluble domain (Figure 4C-D, S6C-D)(*24*). For this, we constructed a combined principal component analysis consisting of all the BioEmu models and 4,000 snapshots taken every 2.5 ns from 5 x 2 μs MD data seeded from either the GlpG structure 2IC8 or 2NRF (see methods for details). This landscape therefore directly compared the BioEmu models and MD simulations. The analysis revealed that, as seen for the BioEmu data alone, a dramatic movement of TM5 predominates the variance, which is represented by eigenvector 1 (Figure 4D). Of note, the distribution of BioEmu models (-2.6 to 0.70) sits entirely within the eigenvector space sampled by the MD, with the 2IC8 MD between -0.49 and 2.5 and 2NRF being between -4.3 and 1.5. Plotting the LG distances clearly demonstrates that this variance is closely connected to LG opening (Figure S6E), with the 2NRF MD simulations sampling both an open and closed conformation; the 2IC8 MD sampling a more closed conformation; and the BioEmu mostly distributed in the middle. This suggests that all states sampled by BioEmu overlap with those sampled with MD. However, both 2IC8- and 2NRF-seeded MD simulations sample states which are more extreme than the BioEmu, relating to a more open or more closed LG. One notable exception is that BioEmu modestly outperforms 2IC8-seeded simulation in sampling open conformations (Figure S6E).

Overall, this demonstrates that BioEmu is able to successfully capture the most salient conformational dynamics of membrane proteins, but may fail to fully explore all possible configurations within the scale of sampling performed in this study.

### BioEmu generates protein ensembles with dramatically reduced resource requirements compared to MD

For proteins which lack solved structures, a common approach to predict conformational ensembles is to use structure prediction tools such as AF2 to generate an input for MD simulations. BioEmu could provide an alternative which reduces the requirement for running costly simulations. To more directly compare these two approaches, we compared sampling achieved by BioEmu alone to a previously published AF2-MD dataset (Figure 5A-B)(*24*). RHBDL1, a human rhomboid intramembrane protease with no identified substrates, was demonstrated to very rarely sample gated open conformations compared to simulations of other rhomboids (∼2% total simulation time above a 1.4 nm gating threshold). Remarkably, BioEmu successfully samples this open conformation, further demonstrating its ability to capture rare states (Figure 5A). Furthermore, we compared the BioEmu prediction for the first 500 ns of each of the 5 simulation runs (2.5 μs total; Figure 5B) in addition to the full 2 μs sampling (10 μs total; Figure 5A). BioEmu outperformed the 500 ns simulations for both the human and bacterial rhomboid and closely matched the 2 μs sampling.

**Figure 5.**
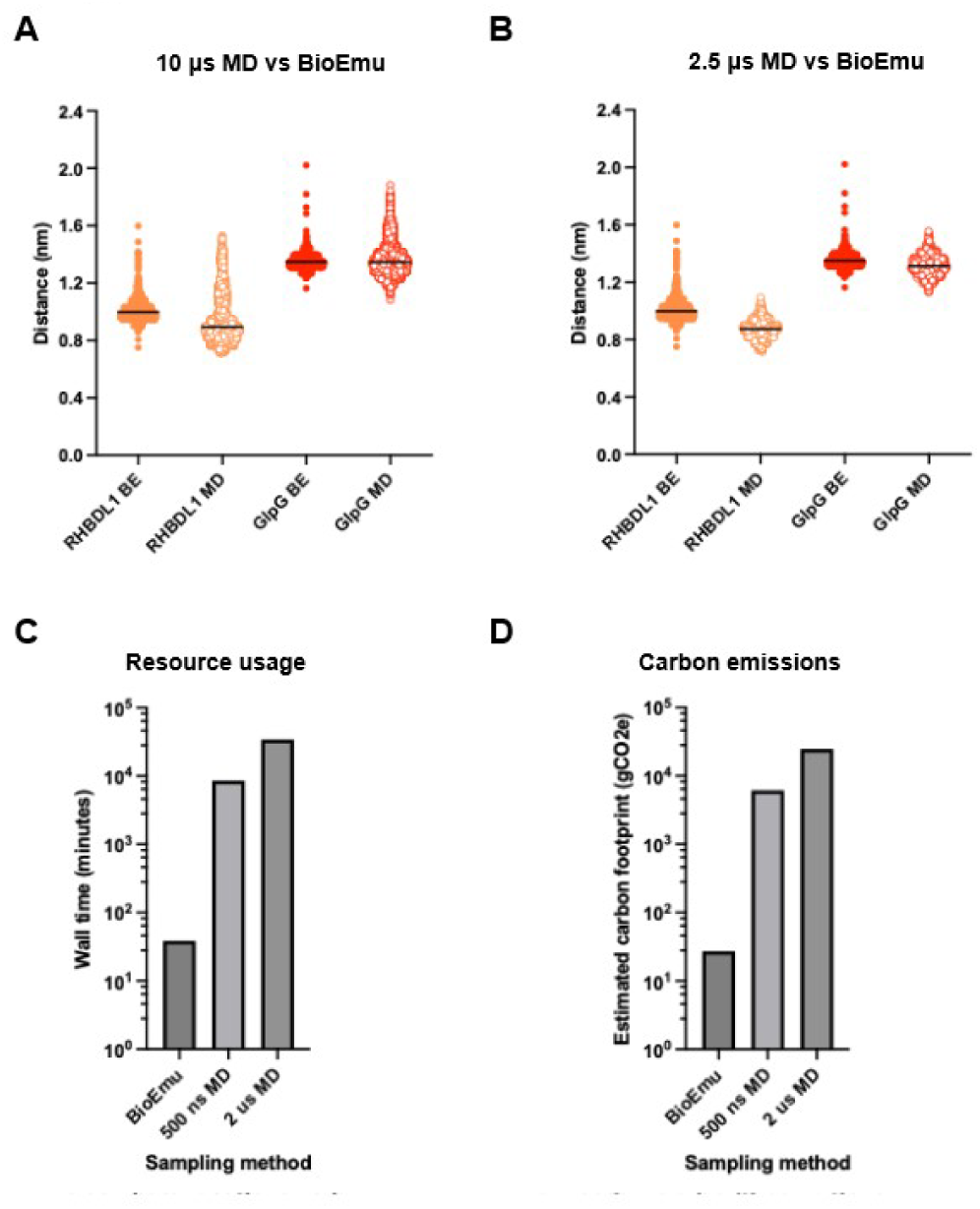
Conformational sampling and resource consumption comparisons for BioEmu vs MD. **(A-B)** Lateral gate distance distribution profiles of human RHBDL1 and *E. coli* GlpG for BioEmu 1000 samples vs 5x 500 ns or 2 μs MD runs. **(C-D)** Estimated resource usage from BioEmu predictions, compared to MD simulations. Calculations were made for GlpG, using Green Algorithms. 1 wall minute = 719.10 mgCO2e for closest the equivalent system and location to those used in this study.

This establishes BioEmu as a valuable and accessible approach for estimating conformational ensembles of membrane proteins which lack solved structures. Moreover, BioEmu uses a fraction of the computational resources. On a single GPU and CPU-powered PC, 1000 BioEmu samples were generated in ∼40 minutes, compared to the >500 hours which would have been required to achieve the 10 μs total simulation time of the same system on the same computer (Figure 5C-D). This equates to a ∼900-fold saving in estimated carbon emissions, or 2.2 tree years (*32*).

## Discussion

In this study, we demonstrate that BioEmu can sample a range of biologically important conformational states for the 276-residue membrane protein GlpG. This provides proof-of-principle for an ability of BioEmu to accurately predict ensembles for proteins outside of its training dataset, expanding the potential of this tool beyond that which was initially proposed(*11*).

In particular, BioEmu was demonstrated to recapitulate dynamics associated with lateral gating, which has widely been accepted as the mechanism by which rhomboids access their transmembrane substrates(*20–24*). Different states associated with gating were identified (Figures 1-3, S4), including several which closely match those of solved GlpG structures(*22, 33–35*). In the open state from this ensemble, outward movement of TM5 coinciding with a lifting of the L5 loop agree with previous reports of a gated open state(*21–23*).

Intermediate states for GlpG were also identified, in which the L5 loop has lifted without significant TM5 movement, and vice versa (Figure S4). An ‘open cap’ structure in which the L5 loop becomes disordered, leading to solvation of the active site, has been previously identified(*36*). Additionally, time-resolved crystallography has led to the proposal of a model under which the L5 loop first lifts, followed by outward movement of TM5. After the substrate enters the active site, the L5 loop then lowers back down to ‘clamp’ the substrate within the scission complex(*37*). The presence of these different states within the BioEmu prediction demonstrates an ability to generate an ensemble that spans the full catalytic cycle of rhomboid proteases even in the absence of substrates, which makes it a powerful tool for interrogating the biological function of a variety of enzymes.

The similarity between predicted and solved states may be due in part to the presence of exprimental GlpG structures in the PDB. This was used for the pretraining of the AlphaFold Evoformer, which forms part of the BioEmu architecture, despite the exclusion of membrane proteins from the training stages of the BioEmu model(*4, 11*). Therefore, future work to interrogate the ability of BioEmu to predict ensembles for membrane proteins without any solved structures should expand this approach to proteins for which structures were added to the PDB *after* the AlphaFold Evoformer was developed.

When BioEmu sampling of rhomboid lateral gate dynamics was directly compared against that of an AF2-MD approach, BioEmu was found to outperform 2.5 μs MD, and achieve very similar results to 10 μs MD (Figure 5)(*24*). Importantly, these results were achieved using ∼900-fold fewer computing resources than the AF2-MD, demonstrating that BioEmu can a broad conformational landscape using a fraction of the computational power of MD.

Despite the impressive performance of BioEmu at capturing lateral gate dynamics, the N-terminal cytoplasmic domain of GlpG was predicted in some models to occupy space which would be filled by the lipid bilayer, making these states physically impossible. Therefore, extra caution should be taken when interpreting BioEmu predictions of transmembrane proteins which also include soluble domains. Despite these erroneous states, physically possible dynamics of the soluble domain of GlpG were also captured, with this region being the most dynamic, as has been previously reported by MD simulations of full-length GlpG generated by AlphaFold(*24*). However, experimental validation of the soluble domain of GlpG in the context of the full protein is currently lacking, with only isolated structures of the soluble domain currently available(*25*).

(Furthermore, a combined principal component analysis of the GlpG BioEmu predictions in the current study, and MD simulations of GlpG structures from a previous study, suggest that BioEmu does not capture all the states sampled in 20 μs of MD seeded from solved structures (Figure 4) (*24*). Additionally, whilst the authors report that BioEmu can accurately predict the equilibrium distribution of protein ensembles, the weighting of the models generated for GlpG by BioEmu differ substantially than that seen in MD simulations (Figures 4 and S6)(*11, 24*). Most notably, only a single population is seen across lateral gate profiles and PCA projections, which does not fit with the expectations for a protein with at least two main conformational states. Additionally, some of the most ‘extreme’ open and closed states seen in MD simulations are not captured by BioEmu. These findings highlight the continued value of MD simulations for applications requiring accurate characterisation of conformational energetics, relative state populations, or the identification of specific rare states. However, it should be noted that neither of the two simulation runs individually samples the full landscape. In fact, the MD distribution from the 2IC8 input model is much more comparable to that of the BioEmu distribution (Figure S6). This reflects a fundamental challenge shared by both approaches: accurate description of protein dynamics depends on sufficient sampling of the relevant conformational space and, for MD simulations, the quality of the initial structural model.

Overall, our results support BioEmu as a rapid and computationally efficient approach for generating biologically relevant conformational ensembles of membrane proteins, whilst highlighting the continued importance of molecular dynamics for accurately characterising conformational energetics.

## Methods

### BioEmu models

BioEmu version 1.2.0 was used to generate 1000 samples per protein (https://github.com/microsoft/bioemu/)(11). As BioEmu filters out unphysical structures, fewer than 1000 models were obtained for each run. The input sequence used and number of samples obtained for each protein is listed in Table 1. The individual frames from the generated trajectory file were extracted as .pdb files using the ‘trjconv’ GROMACS command, with the topology.pdb file as the reference structure(*38*).

**Table 1.** Details of BioEmu models.

| Protein | Input sequence | Number of samples post filtering |
| --- | --- | --- |
| RHBDL1 | All residues of UniProt entry O75783-2 | 881 |
| GlpG | All residues of UniProt entry P09391 | 969 |

BioEmu predictions from this paper are available at the following link: https://osf.io/w5g7c/

### FoldSeek clustering of BioEmu models

BioEmu models were clustered using a local installation of FoldSeek (https://github.com/steineggerlab/foldseek)(30). Default parameters were used, with the exception of the template modelling threshold, which was set to 0.7 for clustering of the full-length GlpG samples. For GlpG ΔN clustering, an index file was created using GROMACS so that a topology file for each sample could be generated, which described only residues 94-276. It should be noted that BioEmu begins its residue numbering at 0, rather than 1, and so these were residues 93-275 using this numbering system. For GlpG ΔN, a template modelling threshold of 0.98 was used to generate clusters.

The clustering of GlpG BioEmu models is outlined in Table 2.

| Full-length GlpG cluster (representative sample) | Samples | GlpG ΔN cluster (representative sample) | Samples |
| --- | --- | --- | --- |
| Cluster 1 (sample 0) | 936 | Cluster 1 (sample 0) | 957 |
| Cluster 2 (sample 26) | 29 | Cluster 2 (sample 707) | 4 |
| Cluster 3 (sample 147) | 3 | Cluster 3 (sample 24) | 1 |
| Cluster 4 (sample 707) | 1 | Cluster 4 (sample 41) | 1 |
|  |  | Cluster 1 (sample 99) | 1 |
|  |  | Cluster 2 (sample 251) | 1 |
|  |  | Cluster 3 (sample 434) | 1 |
|  |  | Cluster 4 (sample 456) | 1 |
|  |  | Cluster 3 (sample 734) | 1 |
|  |  | Cluster 4 (sample 794) | 1 |

### Lipid bilayer plane analysis

To define the plane of the lipid bilayer, the CHARMM-GUI lipid packing coordinate file generated for the AF2 GlpG MD simulations was used as reported previously(*24*). The full-length GlpG BioEmu sample 0 coordinate file was then aligned to this model using PyMOL(*39*), allowing the z co-ordinates within the plane of the lipid bilayer to be identified. The alpha-carbon atoms of the core GlpG fold (residues 94-276) within the BioEmu trajectory file were then fitted against this template, using the fit with ‘rot+trans’ function of the ‘trjconv’ command in GROMACS version 2023.4(*38*). The GROMACS ‘select’ command was then used to identify frames within the BioEmu trajectory file for which any alpha-carbon atoms within the soluble domain (residues 1-81) were within the z-plane of the lipid bilayer. A threshold of 0.5 nm into the plane of the lipid bilayer was applied to define a model as physically unplausible.

### Lateral gate quantification

Lateral gate quantification was performed using GROMACS version 2023.4, as previously described(*24*). Briefly, an average distance across two representative residue pairs across TM2 and TM5 were used. Distances across trajectory files were measured using the ‘mindist’ command.

### RMSD and RMSF analysis

RMSD and RMSF analyses were performed using GROMACS version 2023.4. The trajectory file outputs from BioEmu were used, with the topology.pdb file as the reference structure for RMSF analysis. For RMSD analysis, ‘trimmed’ versions (containing residues 93-270) were created so that numbers of residues matched between all crystal structures and the BioEmu samples. Either the representative model of GlpG ΔN cluster 1 (sample 0), or the crystal structures 2IC8 (chain B due to lack of resolved loop structure in chain A), 2NRF (chain A), 2XOW, or 2IRV (chain A) were used as reference structures.

### Principal component analysis

Principal component analysis was performed using GROMACS version 2023.4. For the GlpG ΔN and combined (GlpG ΔN and MD) analyses, ‘trimmed’ versions (containing residues 93-270) were created so that numbers of residues matched between all crystal structures and the BioEmu samples. GROMACS ‘covar’ and ‘anaeig’ were used to construct covariance matrices and analyses covariance matrices.

### Resource usage calculations

To calculate equivalent wall time comparisons for the generation of BioEmu predictions versus MD simulations, the time required for BioEmu to generate all 1000 predictions of full-length GlpG in the present study was recorded. A single-node GPU- and CPU-powered computer was used for these experiments. A short, 5-minute test simulation of the post-equilibration GlpG model system inside a lipid bilayer from a previous study(*24*) was run on the same system. The hours/s from the log file from this were used to extrapolate the total time required to sample 10 μs on the same node.

CO_2_ emissions were estimated by using the wall times as input for Green Algorithms, selecting the closest GPU and CPU models available to those which were used in this study(*32*).

### Data visualisation

Models of proteins were visualised and structural alignments (with the Matchmaker tool) performed using ChimeraX(*40*). Graphs were generated using GraphPad Prism 10.2.0(*41*).

### Data and software availability

All BioEmu models from this study are available for download from https://osf.io/w5g7c/. MD datasets are available from https://osf.io/n6348/. Analysis scripts are available on GitHub (https://github.com/bryonyclifton/AnalysisScripts)

---

**Figure S1.**
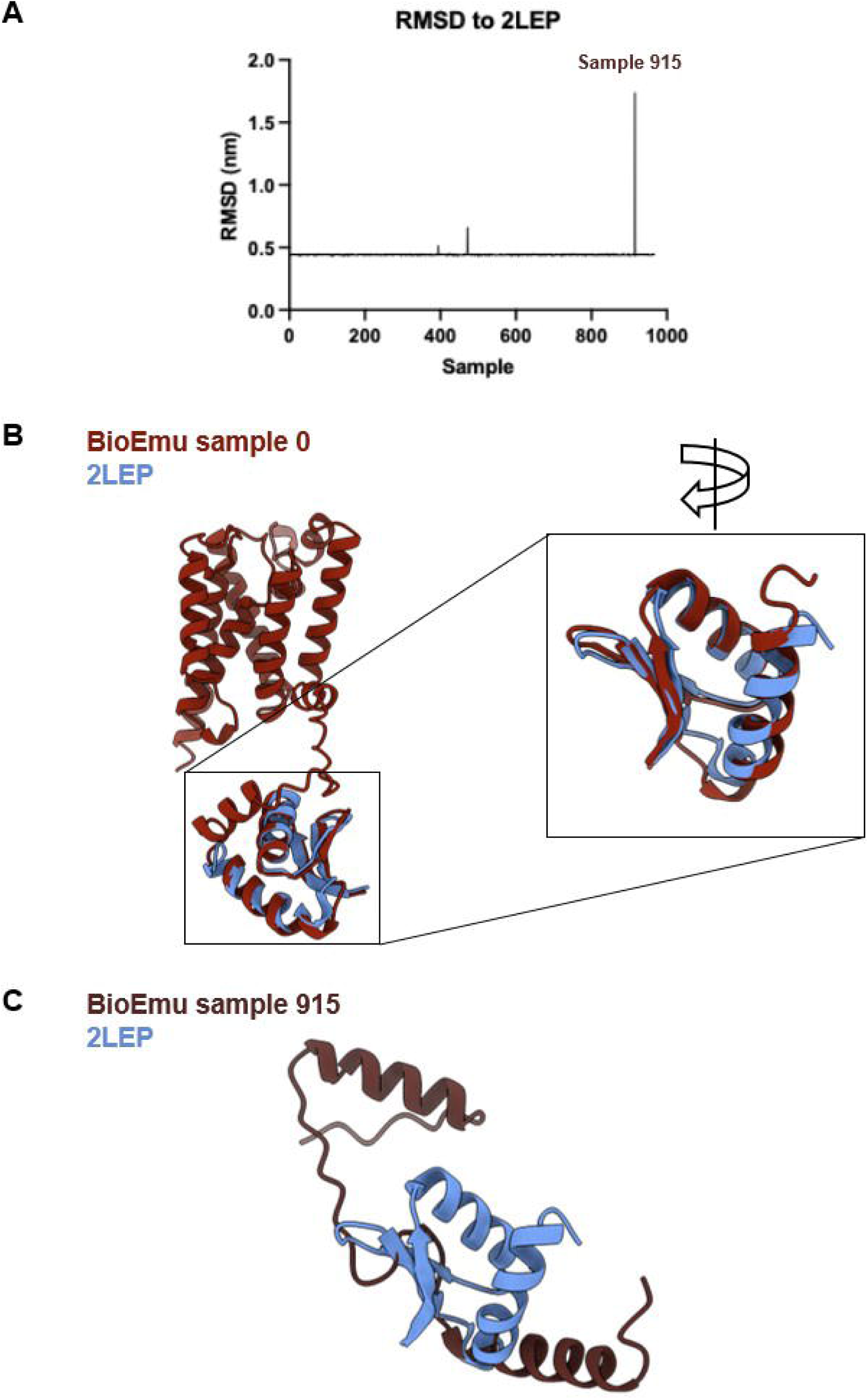

**Figure S2.**
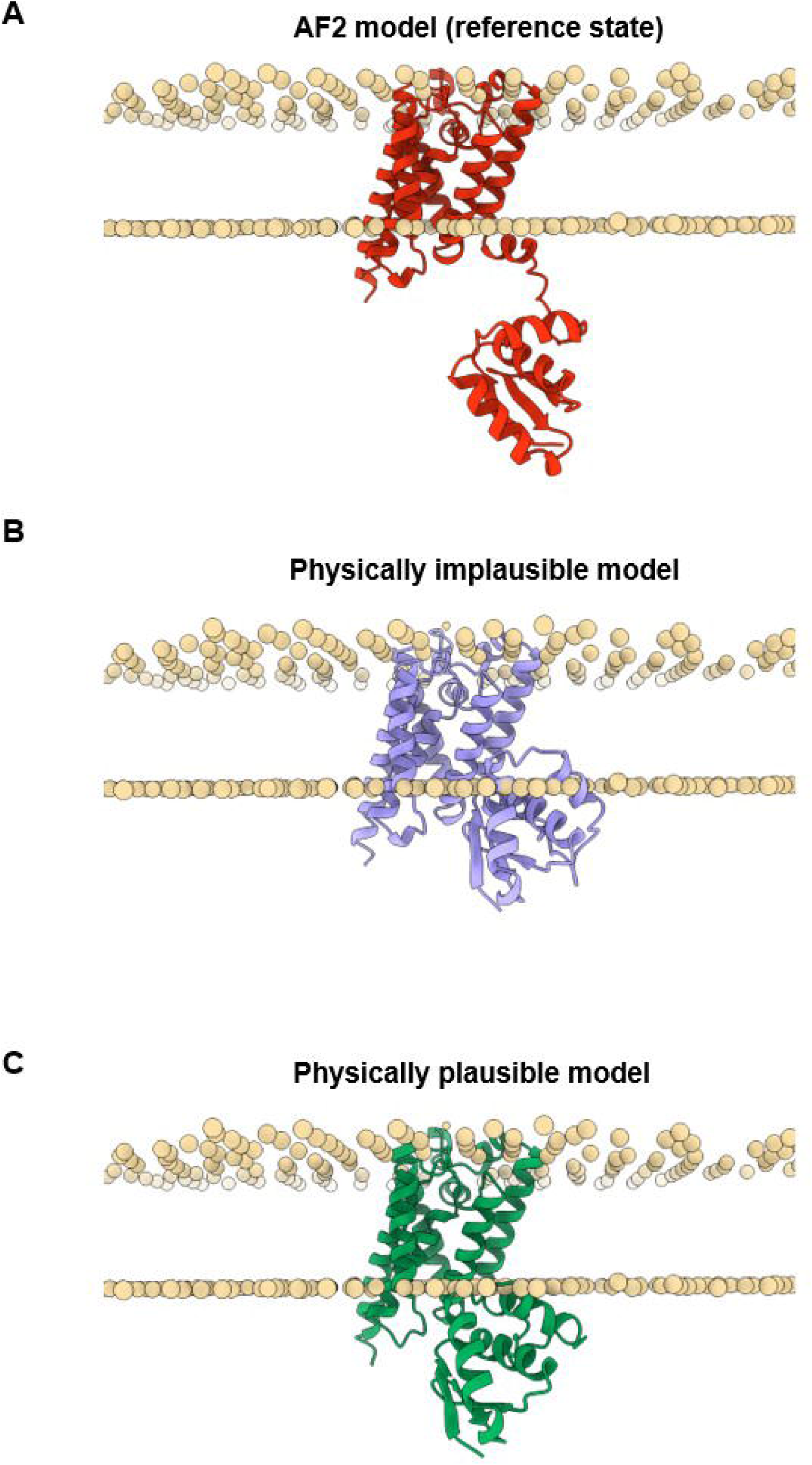

**Figure S3.**
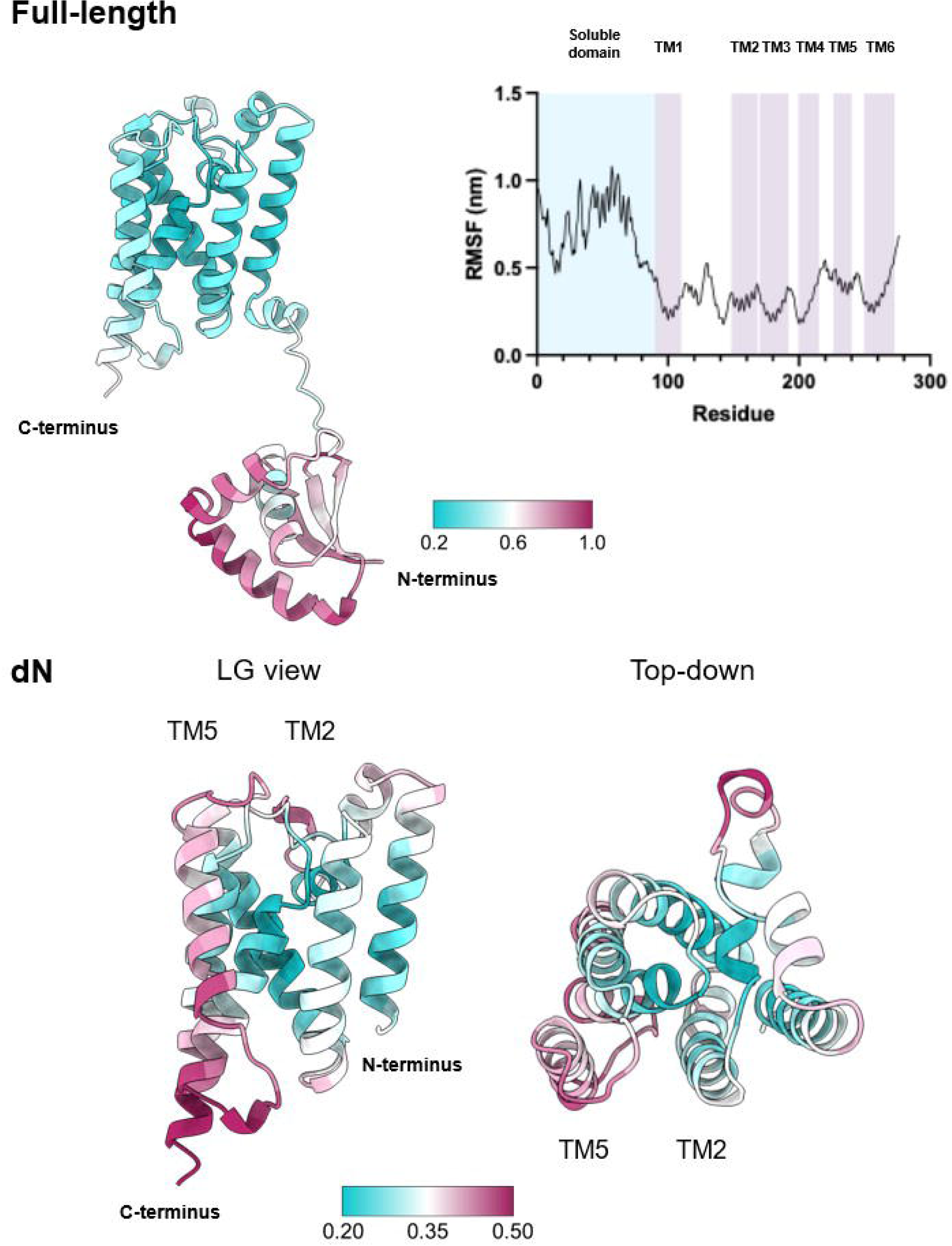

**Figure S4.**
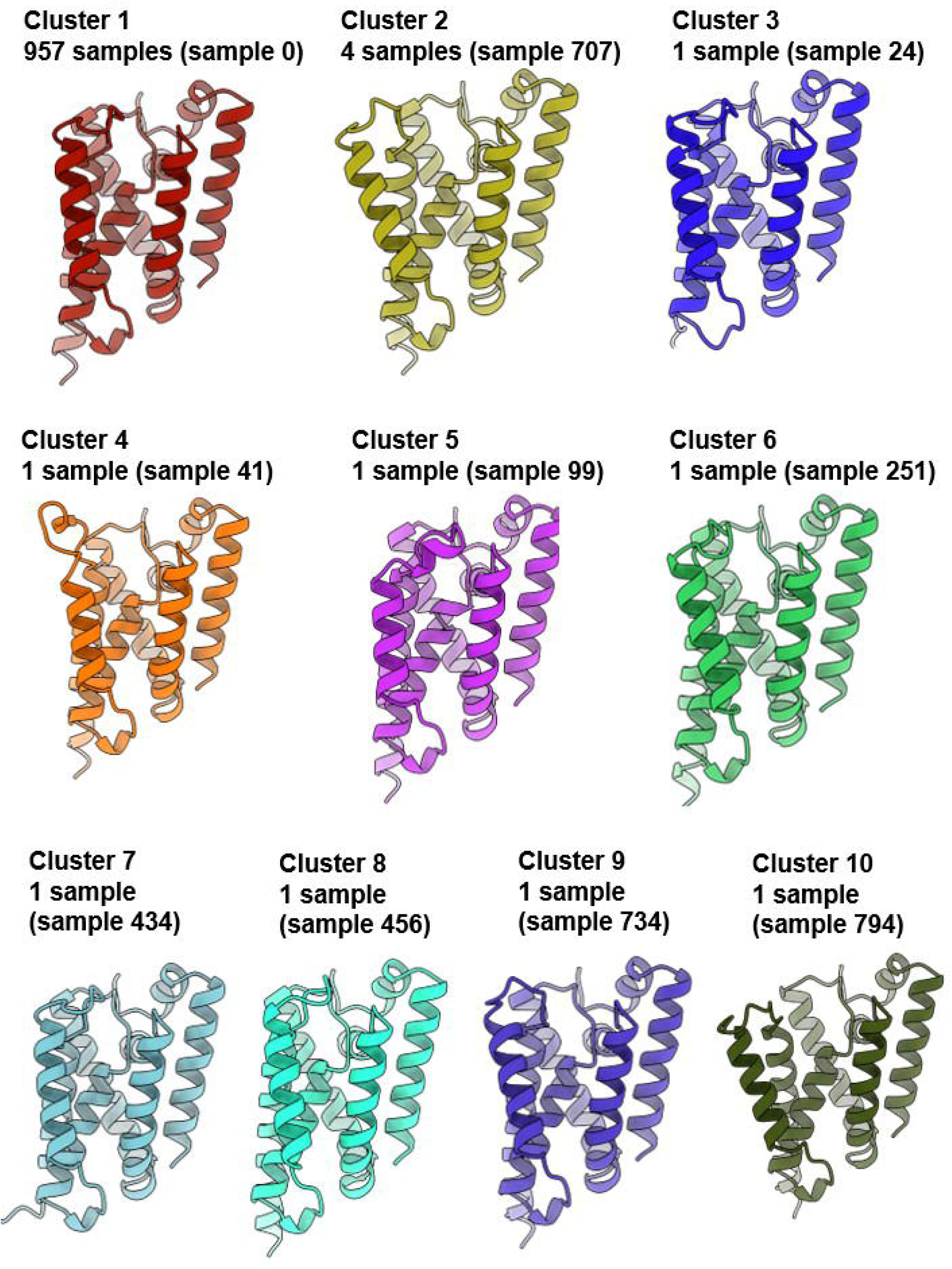

**Figure S5.**
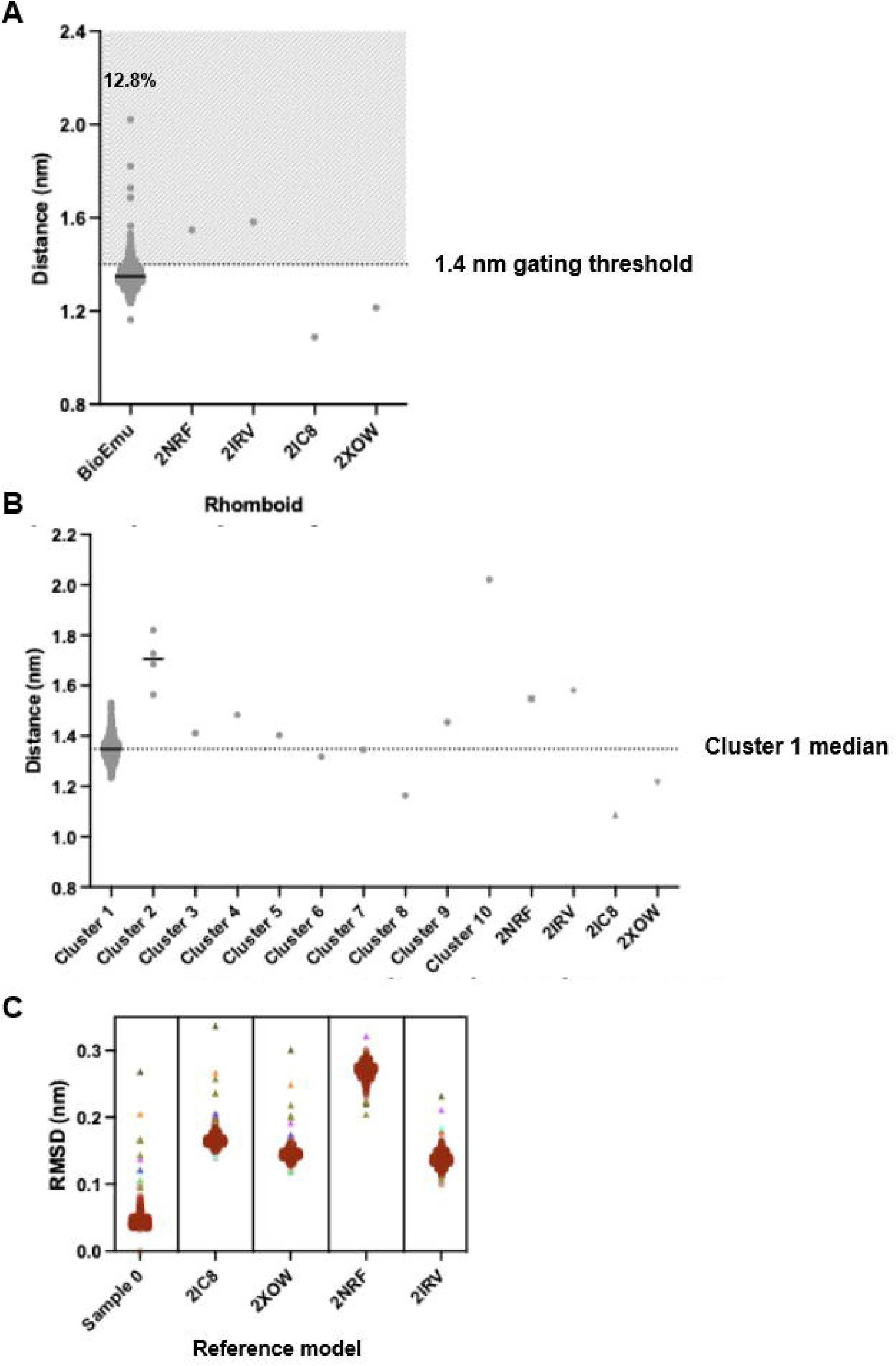

**Figure S6.**
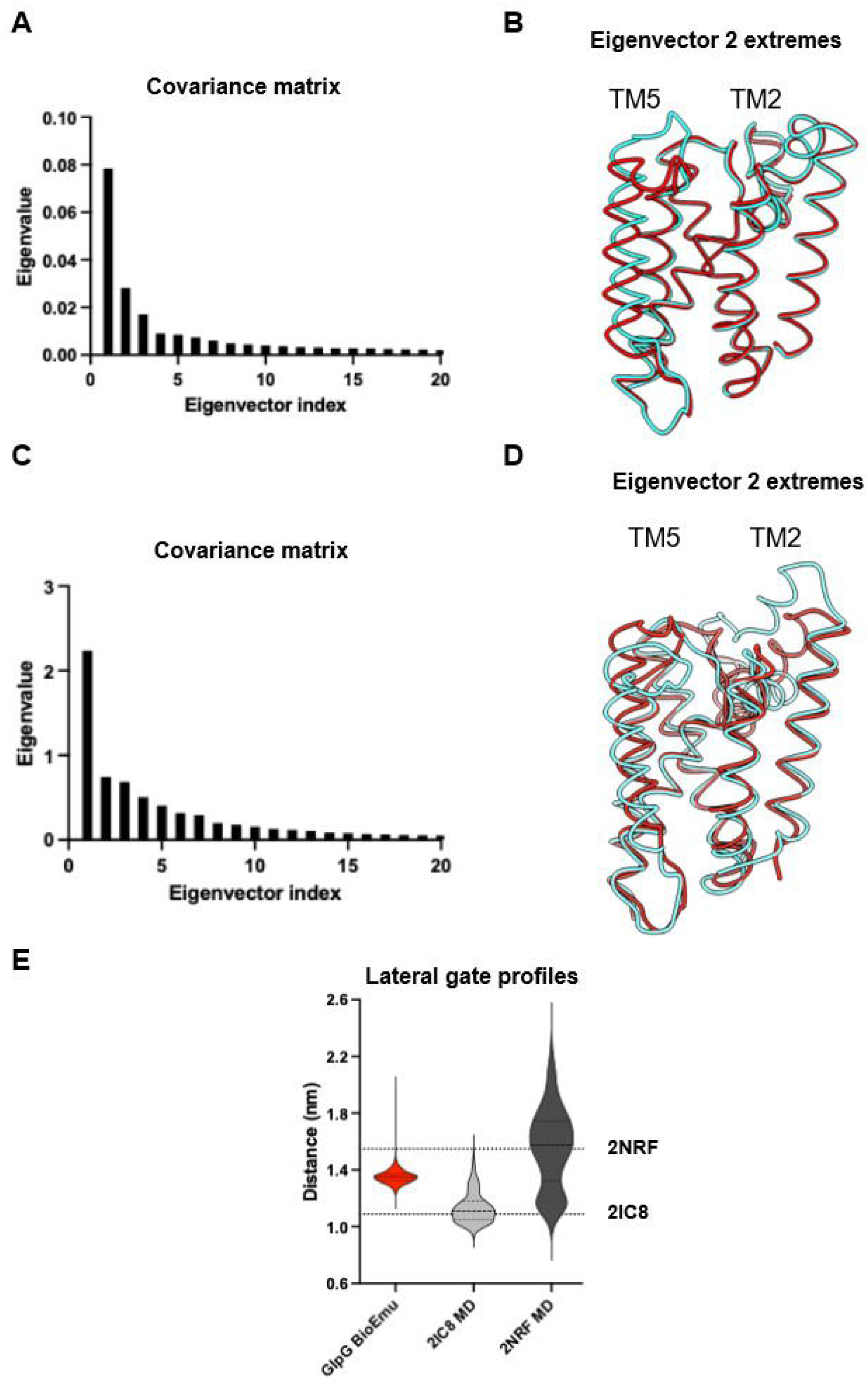

**Figure S7.**
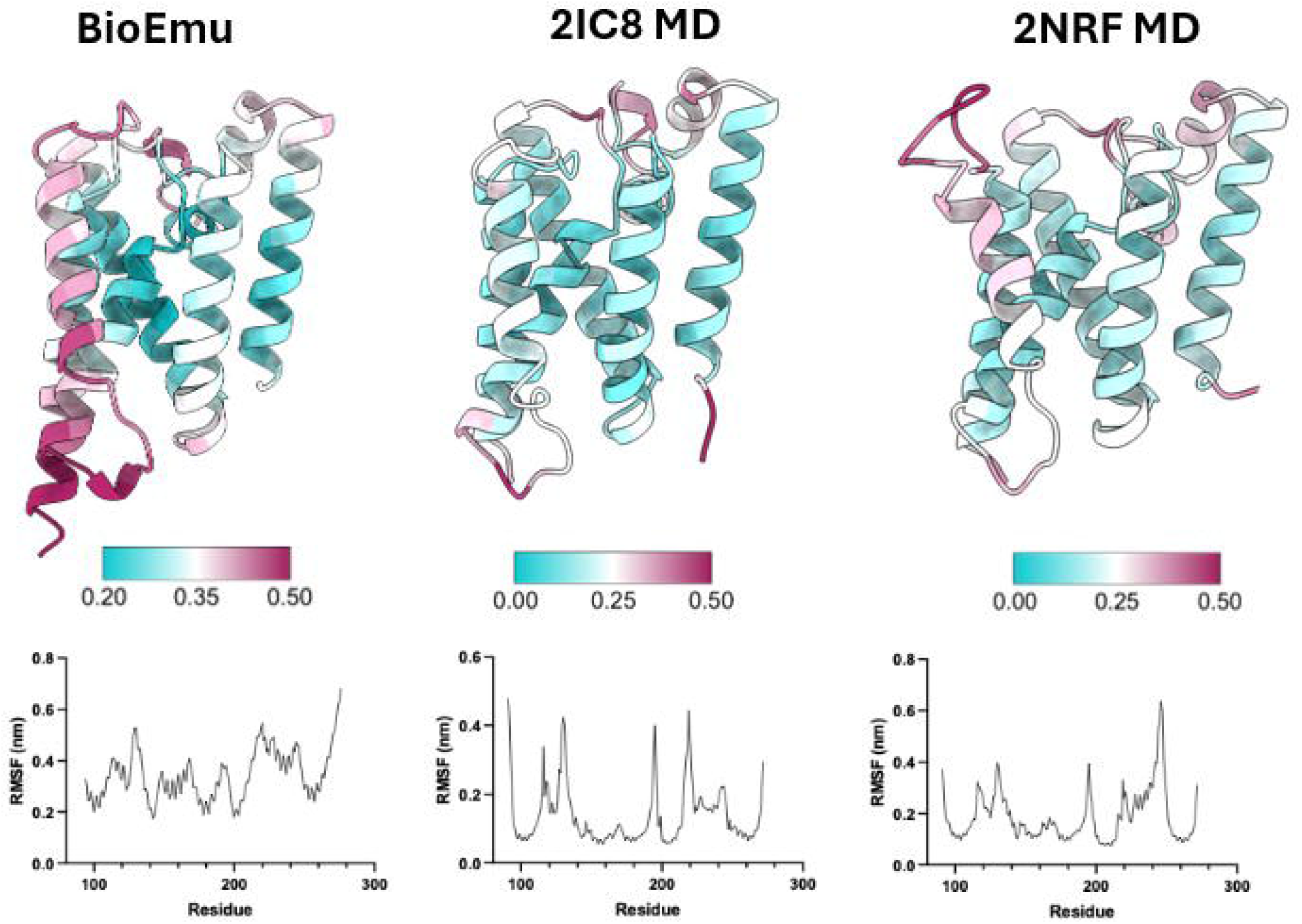

## References

1. J. Berriman, N. Unwin, Analysis of transient structures by cryo-microscopy combined with rapid mixing of spray droplets. Ultramicroscopy 56, 241–252 (1994).

2. B. Chen et al., Structural dynamics of ribosome subunit association studied by mixing-spraying time-resolved cryogenic electron microscopy. Structure 23, 1097–1105 (2015).

3. M. Jelokhani-Niaraki, Membrane Proteins: Structure, Function and Motion. Int J Mol Sci 24, (2022).

4. J. Jumper et al., Highly accurate protein structure prediction with AlphaFold. Nature 596, 583–589 (2021).

5. J. Abramson et al., Accurate structure prediction of biomolecular interactions with AlphaFold 3. Nature 630, 493–500 (2024).

6. T. Hegedűs, M. Geisler, G. L. Lukács, B. Farkas, Ins and outs of AlphaFold2 transmembrane protein structure predictions. Cell Mol Life Sci 79, 73 (2022).

7. M. A. Jambrich, G. E. Tusnady, L. Dobson, How AlphaFold2 shaped the structural coverage of the human transmembrane proteome. Sci Rep 13, 20283 (2023).

8. G. Monteiro da Silva, J. Y. Cui, D. C. Dalgarno, G. P. Lisi, B. M. Rubenstein, High-throughput prediction of protein conformational distributions with subsampled AlphaFold2. Nat Commun 15, 2464 (2024).

9. D. Sala, F. Engelberger, H. S. McHaourab, J. Meiler, Modeling conformational states of proteins with AlphaFold. Curr Opin Struct Biol 81, 102645 (2023).

10. C. Upex, T. Osborne, G. Biglino, J. Hancox, R. A. Corey, Integrating AI and molecular modeling for structural prediction of a closed state of the hERG channel. bioRxiv, 2026.2004.2020.719540 (2026).

11. S. Lewis et al., Scalable emulation of protein equilibrium ensembles with generative deep learning. Science, eadv9817 (2025).

12. J. Zha et al., Assessing the Performance of BioEmu in Understanding Protein Dynamics. Int J Mol Sci 27, (2026).

13. H. Sawczyc, S. Kosteletos, A. Lange, Death, taxes, and rhomboids: Understanding the ubiquitous roles of the rhomboid protein superfamily. J Biol Chem 301, 110699 (2025).

14. J. R. Lee, S. Urban, C. F. Garvey, M. Freeman, Regulated intracellular ligand transport and proteolysis control EGF signal activation in Drosophila. Cell 107, 161–171 (2001).

15. M. K. Lemberg, M. Freeman, Cutting proteins within lipid bilayers: rhomboid structure and mechanism. Mol Cell 28, 930–940 (2007).

16. M. Freeman, The rhomboid-like superfamily: molecular mechanisms and biological roles. Annu Rev Cell Dev Biol 30, 235–254 (2014).

17. V. L. Lastun, A. G. Grieve, M. Freeman, Substrates and physiological functions of secretase rhomboid proteases. Semin Cell Dev Biol 60, 10–18 (2016).

18. A. G. Grieve et al., Conformational surveillance of Orai1 by a rhomboid intramembrane protease prevents inappropriate CRAC channel activation. Mol Cell 81, 4784–4798.e4787 (2021).

19. S. W. Dickey, R. P. Baker, S. Cho, S. Urban, Proteolysis inside the membrane is a rate-governed reaction not driven by substrate affinity. Cell 155, 1270–1281 (2013).

20. C. Bohg et al., The opening dynamics of the lateral gate regulates the activity of rhomboid proteases. Sci Adv 9, eadh3858 (2023).

21. R. P. Baker, K. Young, L. Feng, Y. Shi, S. Urban, Enzymatic analysis of a rhomboid intramembrane protease implicates transmembrane helix 5 as the lateral substrate gate. Proc Natl Acad Sci U S A 104, 8257–8262 (2007).

22. Z. Wu et al., Structural analysis of a rhomboid family intramembrane protease reveals a gating mechanism for substrate entry. Nat Struct Mol Biol 13, 1084–1091 (2006).

23. S. Urban, R. P. Baker, In vivo analysis reveals substrate-gating mutants of a rhomboid intramembrane protease display increased activity in living cells. Biol Chem 389, 1107–1115 (2008).

24. B. R. Clifton, R. A. Corey, A. G. Grieve, Structural and energetic insights into human rhomboid proteases reveal a unique lateral gating mechanism for orphan family members. bioRxiv, 2025.2011.2021.689725 (2026).

25. A. R. Sherratt, D. R. Blais, H. Ghasriani, J. P. Pezacki, N. K. Goto, Activity-based protein profiling of the Escherichia coli GlpG rhomboid protein delineates the catalytic core. Biochemistry 51, 7794–7803 (2012).

26. Y. Wang, Y. Zhang, Y. Ha, Crystal structure of a rhomboid family intramembrane protease. Nature 444, 179–180 (2006).

27. R. P. Baker, S. Urban, Cytosolic extensions directly regulate a rhomboid protease by modulating substrate gating. Nature 523, 101–105 (2015).

28. J. J. Lim et al., Structural insights into the interaction of p97 N-terminus domain and VBM in rhomboid protease, RHBDL4. Biochemical Journal 473, 2863–2880 (2016).

29. M. Jin et al., Structural characterization of metal binding in human tyrosylprotein sulfotransferase 2, TPST2. Sci Rep 16, 6066 (2026).

30. M. van Kempen et al., Fast and accurate protein structure search with Foldseek. Nat Biotechnol 42, 243–246 (2024).

31. M. A. Lomize, I. D. Pogozheva, H. Joo, H. I. Mosberg, A. L. Lomize, OPM database and PPM web server: resources for positioning of proteins in membranes. Nucleic Acids Research 40, D370–D376 (2012).

32. L. Lannelongue, J. Grealey, M. Inouye, Green Algorithms: Quantifying the Carbon Footprint of Computation. Adv Sci (Weinh) 8, 2100707 (2021).

33. C. Shi et al., Structure and Dynamics of the Rhomboid Protease GlpG in Liposomes Studied by Solid-State NMR. J Am Chem Soc 141, 17314–17321 (2019).

34. A. Ben-Shem, D. Fass, E. Bibi, Structural basis for intramembrane proteolysis by rhomboid serine proteases. Proc Natl Acad Sci U S A 104, 462–466 (2007).

35. K. R. Vinothkumar et al., The structural basis for catalysis and substrate specificity of a rhomboid protease. Embo j 29, 3797–3809 (2010).

36. Y. Wang, Y. Ha, Open-cap conformation of intramembrane protease GlpG. Proc Natl Acad Sci U S A 104, 2098–2102 (2007).

37. S. Cho, R. P. Baker, M. Ji, S. Urban, Ten catalytic snapshots of rhomboid intramembrane proteolysis from gate opening to peptide release. Nat Struct Mol Biol 26, 910–918 (2019).

38. D. Van Der Spoel et al., GROMACS: fast, flexible, and free. J Comput Chem 26, 1701–1718 (2005).

39. L. Schrödinger, W. DeLano. (2022).

40. E. F. Pettersen et al., UCSF ChimeraX: Structure visualization for researchers, educators, and developers. Protein Sci 30, 70–82 (2021).

41. GraphPad Prism Version 10 GraphPad Software, San Diego, California USA, www.graphpad.com (software) (2023).

