## Supplementary figures for "Benchmarking AI-generated structural ensembles of membrane proteins against physics-based modelling"

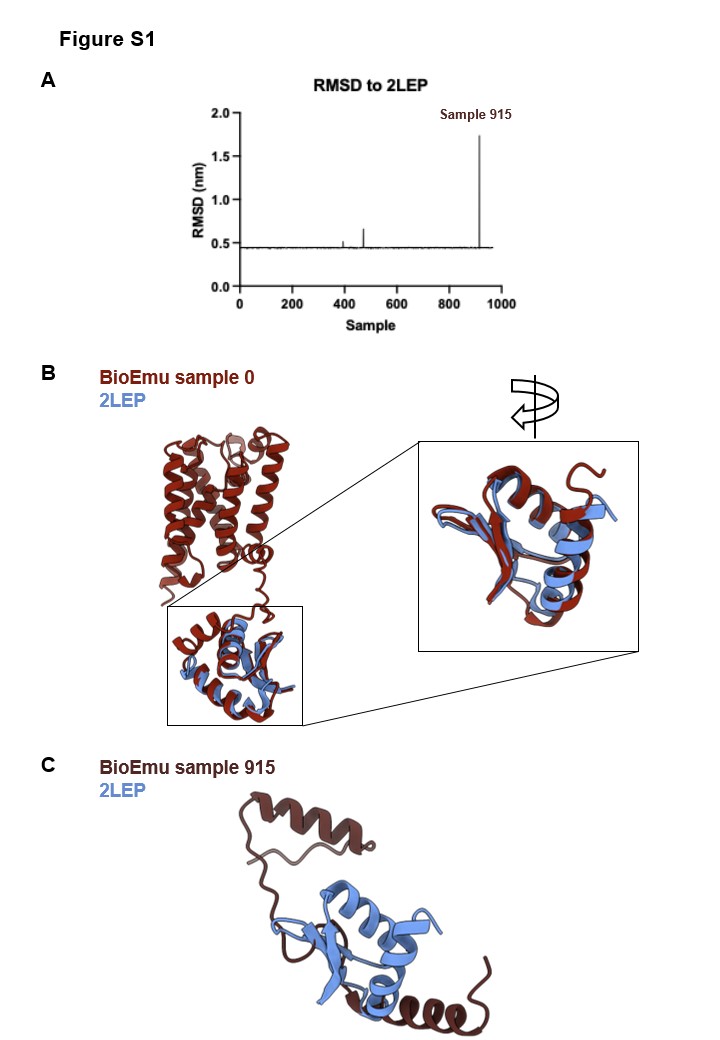

**Figure S1. (A)** RMSD analysis of the GlpG soluble domain from BioEmu samples against an NMR structure (PDB entry 2LEP; model 1) of the equivalent residues(*25*). **(B)** Structural overlays of BioEmu sample 0 with 2LEP. **(C)** Structural overlay of the BioEmu sample 915 soluble domain with 2LEP.

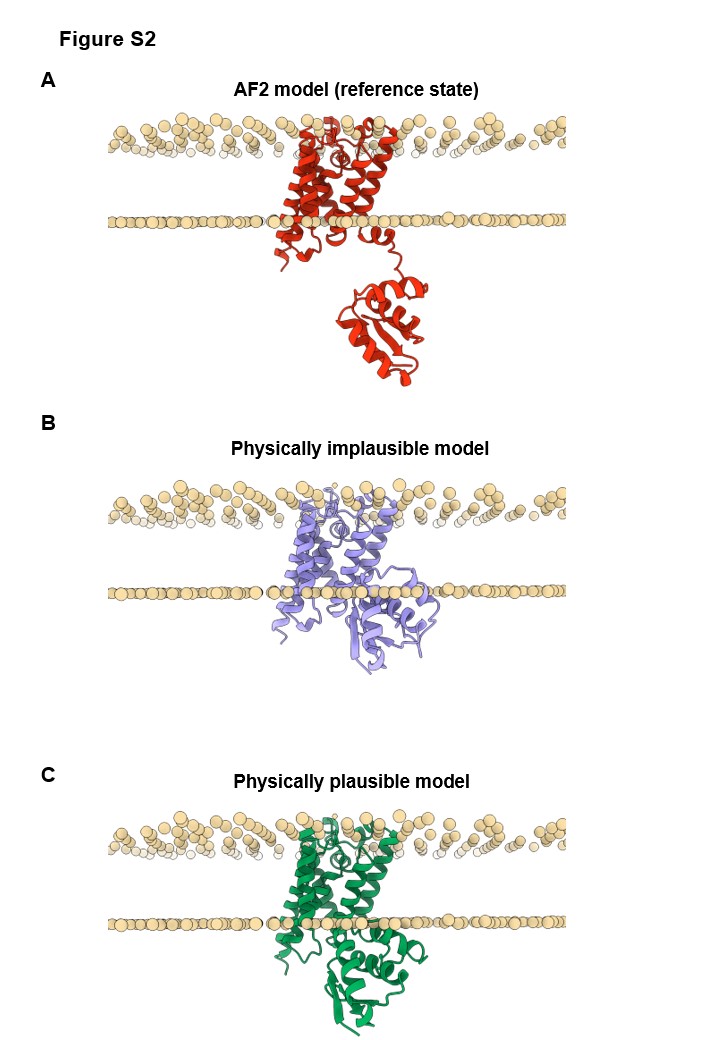

**Figure S2. (A)** Cartoon representation of a GlpG AF2 model embedded in a lipid bilayer. POPC lipid head groups are shown as spheres. **(B)** An example of a BioEmu GlpG model (sample 20, cluster 1) which was considered physically implausible due to membrane clashes (>0.5 nm penetration). **(C)** An example of a GlpG model (sample 49, cluster 1) which was considered plausible, despite minor overlap with the lipid bilayer (< 0.5 nm penetration).

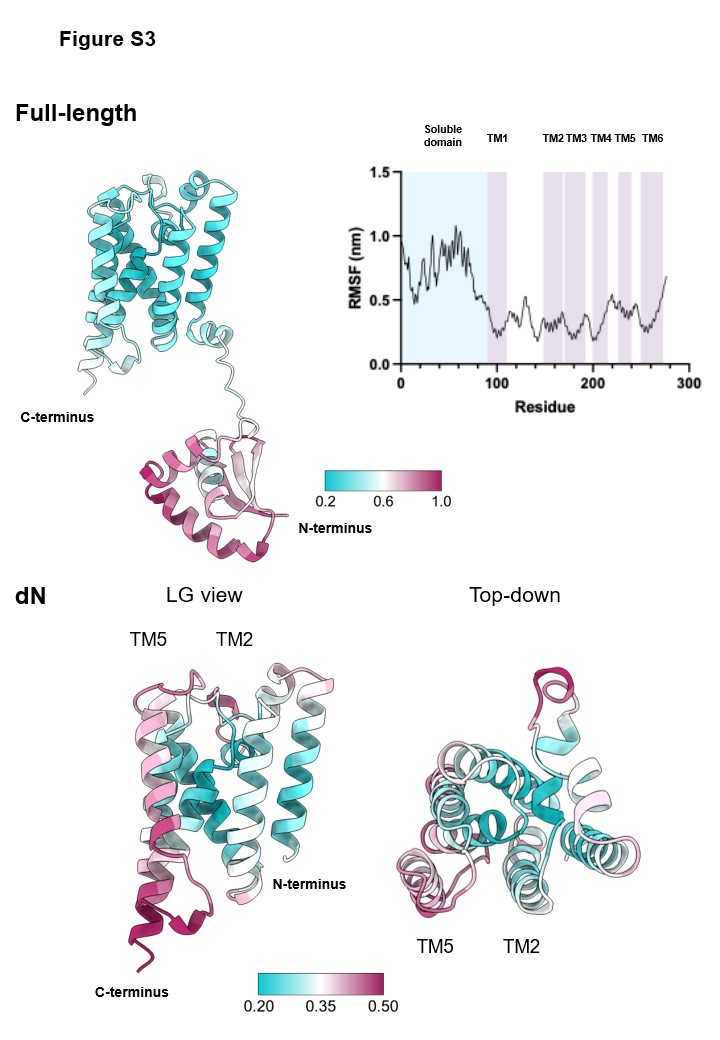

**Figure S3.** RMSF analysis of BioEmu predictions of GlpG. RMSF values are projected onto cartoon depictions of full-length GlpG (top) or GlpG ΔN (bottom).

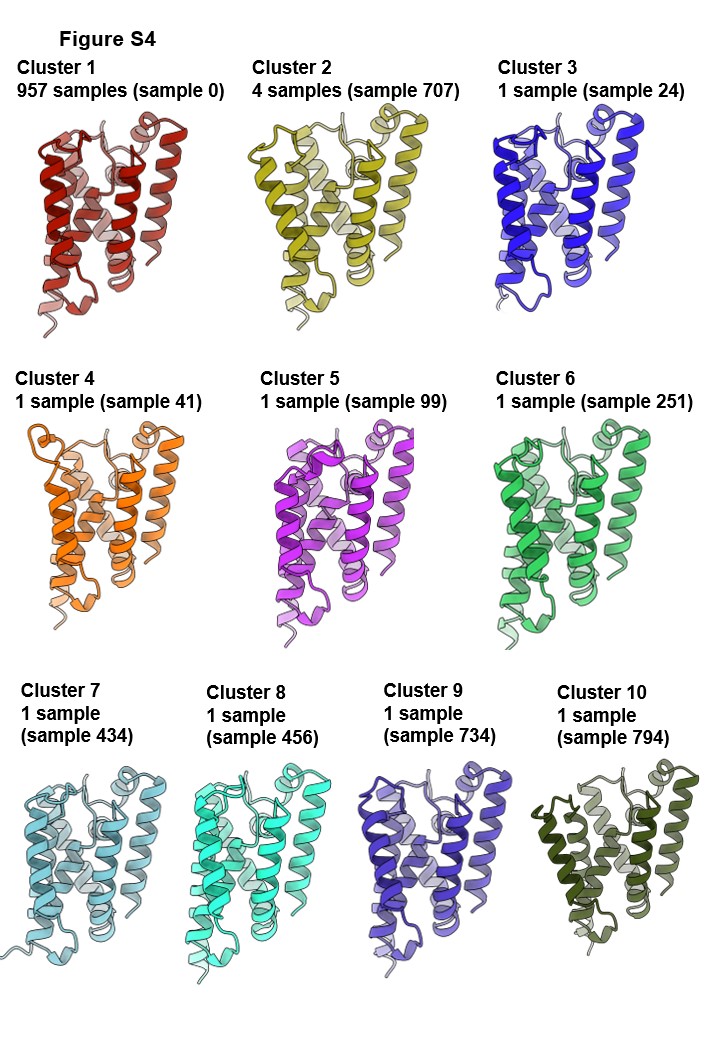

**Figure S4.** Atoms from residues Met1 to Ala93 were removed to form GlpG ΔN. FoldSeek clustering was then applied, using a template modelling threshold of 0.98. The representative model for each cluster is depicted by a cartoon representation and the number of models in each cluster is indicated.

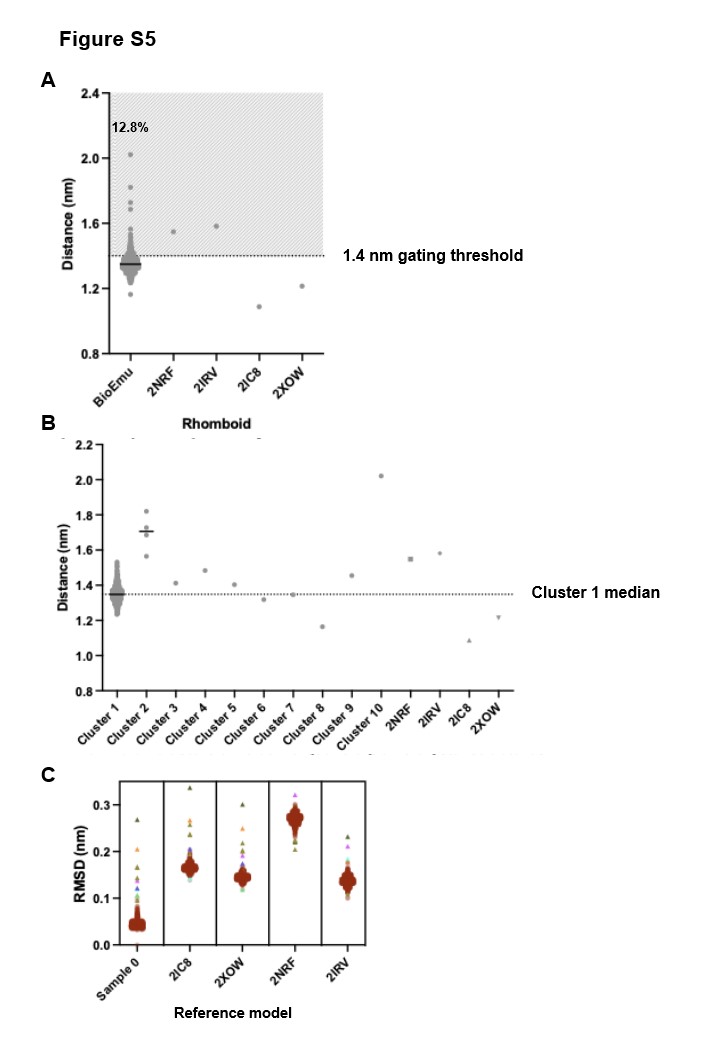

**Figure S5. (A)** The lateral gate distance for each BioEmu sample was quantified and plotted against that of PDB structures 2NRF, 2IRV, 2IC8 and 2XOW. Each point is a model. Line = median. The dashed line indicates the approximate rhomboid gating threshold of c.a. 1.4 nm. 12.8% of models feature a lateral gate above this measurement. **(B)** The equivalent quantification as in (A) was performed and each cluster is shown separately along the x-axis. The dashed line indicates that lateral gate width of the median cluster 1 model. **(C)** RMSD analysis of each BioEmu sample against the reference indicated along the x-axis. All cluster 1 samples are shown as translucent red circles. All other clusters are opaque triangles, with colours matching those in Figure S2.

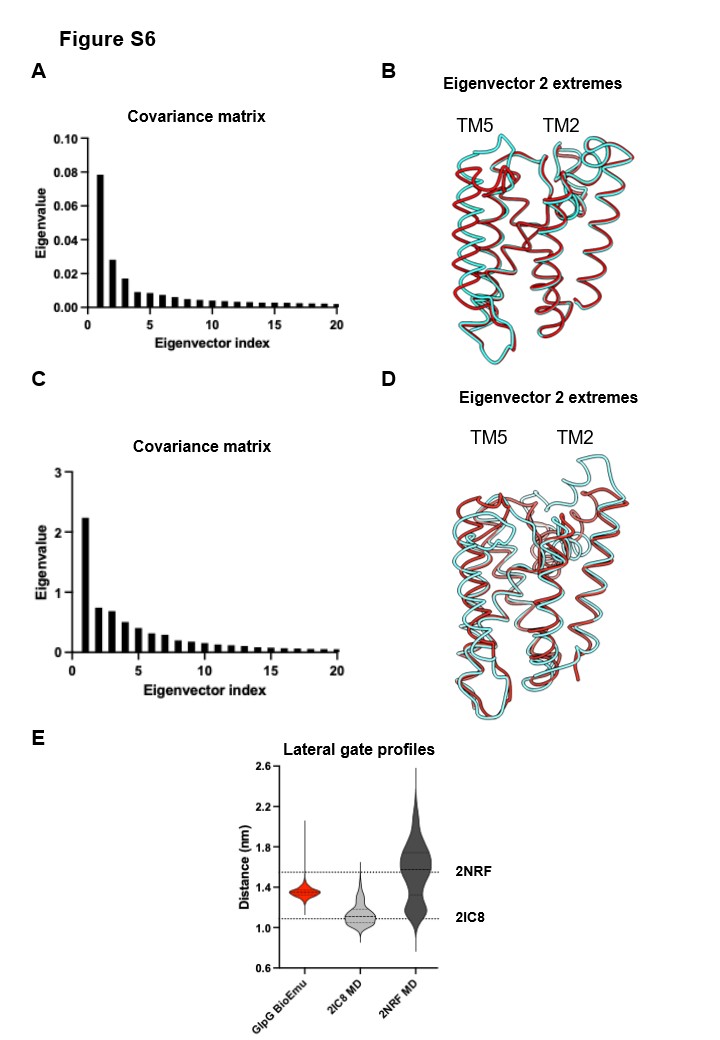

**Figure S6. (A)** Covariance matrix for GlpG trimmed BioEmu PCA. Eigenvectors 1 and 2 contribute 27.6% and 9.91% total variance respectively. **(B)** Overlays of eigenvector 2 extremes for GlpG trimmed BioEmu PCA. **(C)** Covariance matrix for combined BioEmu and MD GlpG PCA. Eigenvectors 1 and 2 contribute 28.9% and 9.58% total variance respectively. **(D)** Overlays of eigenvector 2 extremes for MD and BioEmu combined GlpG PCA. **(E)** Lateral gate distance profiles for GlpG BioEmu vs MD simulations seeded with either 2IC8 or 2NRF(*24*). Dashed lines indicate the lateral gate distance in crystal structures.

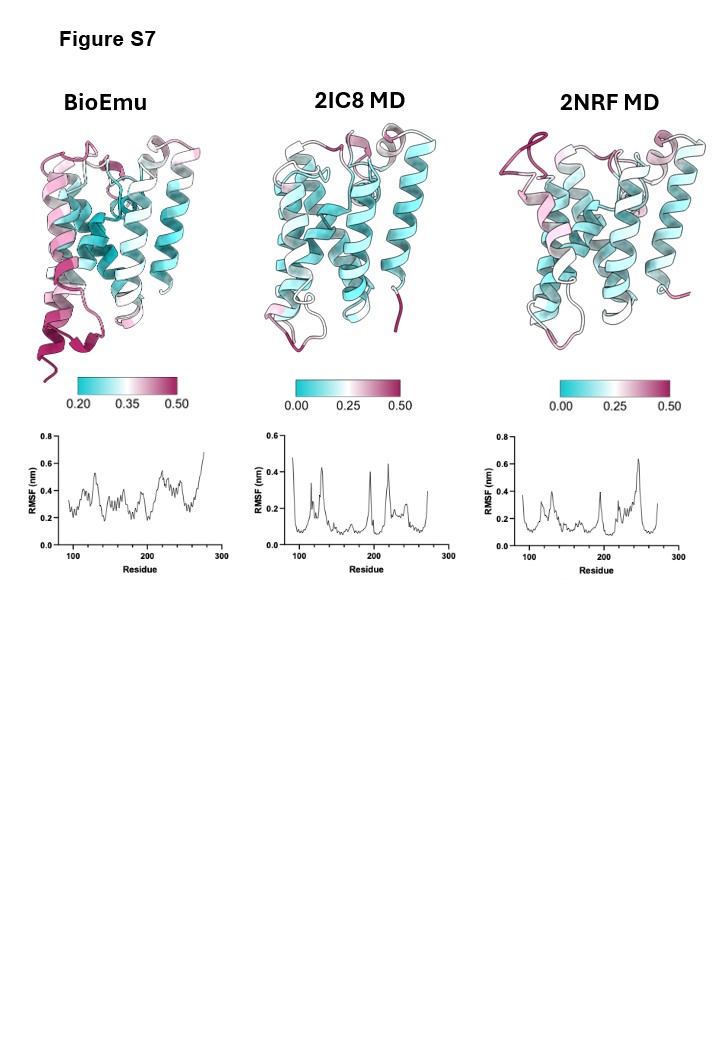

**Figure S7.** RMSF comparisons for GlpG across BioEmu, 2IC8 MD and 2NRF MD. RMSF values (bottom) are projected onto cartoon depictions of GlpG (top).
